# ABPP-HT - high-throughput activity-based profiling of deubiquitylating enzyme inhibitors in a cellular context

**DOI:** 10.1101/2020.12.10.419499

**Authors:** Hannah Jones, Raphael Heilig, Roman Fischer, Benedikt M Kessler, Adan Pinto-Fernandez

## Abstract

The potency and selectivity of a small molecule inhibitor are key parameters to assess during the early stages of drug discovery. In particular, it is very informative for characterizing compounds in a relevant cellular context in order to reveal potential off-target effects and drug efficacy. Activity-based probes (ABPs) are valuable tools for that purpose, however, obtaining cellular target engagement data in a high-throughput format has been particularly challenging. Here, we describe a new methodology named ABPP-HT (high-throughput-compatible activity-based protein profiling), implementing a semi-automated proteomic sample preparation workflow that increases the throughput capabilities of the classical ABPP workflow approximately ten times while preserving its enzyme profiling characteristics. Using a panel of deubiquitylating enzyme (DUB) inhibitors, we demonstrate the feasibility of ABPP-HT to provide compound selectivity profiles of endogenous DUBs in a cellular context at a fraction of time as compared to previous methodologies.

## 1 Introduction

Activity-based probes (ABPs) can assess enzyme activity and inhibition within a cellular environment, thereby providing a considerable advantage over classical biochemical and enzyme assays. ABPs typically consist of a reactive group that binds to the active site of an enzyme, mostly in a covalent fashion, a specific binding group/linker to aid target binding/prevent steric hindrance, and a reporter tag for fluorescence or affinity (Chen et al., 2017; Chakrabarty et al., 2019; Deng et al., 2020). ABPs with an electrophilic reactive group can be applied to multiple enzymes including serine hydrolases, kinases, metalloproteases and cysteine proteases (Niphakis and Cravatt, 2014). Where a nucleophilic active site does not exist, photo-affinity probes can be applied instead (Niphakis and Cravatt, 2014; Mathur et al., 2020). A very informative application of these probes is the activity-based protein profiling (ABPP), combining labeling with ABPs, immunoaffinity purification, and mass spectrometry (IAP-MS) to analyze the probe interactome (Benns et al., 2021). ABPP has been used to study the potency and selectivity of small molecule inhibitors in an unbiased manner, in a cellular matrix (Nguyen et al., 2017; Wang et al., 2018).

Profiling the ubiquitin conjugating activity of E3 ubiquitin ligases and deubiquitylating enzymes (DUBs) is of significant interest due to post-translational protein ubiquitination regulating numerous cellular pathways. These include protein degradation, localization or controlling function (Hershko and Ciechanover, 1992; Mukhopadhyay and Riezman, 2007). Dysregulation of ubiquitination has been linked to several pathologies including cancers and neurodegenerative diseases (Popovic et al., 2014). Consequently, various E3 ligases and DUBs are being evaluated as potential drug targets for either protein targeting chimeras (PROTACs) or inhibitors, respectively, due to their proven ability to specifically target cellular protein homeostasis (Huang and Dixit, 2016; Lai and Crews, 2017; Harrigan et al., 2018).

DUBs offer a mechanistic entry point for probe targeting as the majority are cysteine proteases, with a smaller subsection (< 15 %) functioning as metalloproteases (Komander et al., 2009; Clague et al., 2019). The activity of thiol protease DUBs can be ascertained via covalent attachment of an electrophile to their nucleophilic active site within the catalytic domain. This was first achieved by replacing the C-terminal glycine 76 of ubiquitin with glycyl vinyl sulfone (Borodovsky et al., 2001), and further developed towards a panel of seven different Ub probes with different electrophilic warheads (Borodovsky et al., 2002). These studies have provided the framework for expanding the ubiquitin-based probe concept regarding synthesis (El Oualid et al., 2010) and chemical capture (Hewings et al., 2017), but also targeting metallo-DUBs (Hameed et al., 2019) and E3 ligases (Mulder et al., 2016; Pao et al., 2016).

The inclusion of an affinity tag such as hemagglutinin (HA), FLAG, biotin, etc. to the N-terminus of a ubiquitin-based probe allows for DUB enrichment by immunoprecipitation (IP). Subsequent analysis by liquid chromatography tandem mass spectrometry (LC-MS/MS) can identify and quantify cellular active DUBs bound to the probe. This method can be used in conjunction with a DUB inhibitor to identify inhibitor potency and cross-reactivity. Any DUBs that react with the inhibitor will not bind to the probe as efficiently and will be reduced in the immunoprecipitated sample when compared to a control. This method was successfully applied to demonstrate the selectivity of a USP7 inhibitor, with a 2-bromoethylamine warhead probe (HA-Ub-Br2). In this case, a panel of 22 DUBs were quantified (Turnbull et al., 2017). The ABPP assay was used also to assess cellular target engagement and DUB selectivity in crude cell extracts for small molecule inhibitors against USP9X (Clancy et al., 2020), and USP28 (Ruiz et al., 2020).

Without fractionation the number of DUBs immunoprecipitated and quantified by MS with a propargylamine (PA) warhead is around 30-40 (Altun et al., 2011; Ekkebus et al., 2013). To improve this methodology, we have recently combined this probe (Ub-PA) with sample fractionation and 74 DUBs in MCF-7 breast cancer cells were quantified. For comparison, the transcriptomics analysis of the same cells identified a very similar number of (78) DUB mRNAs (Pinto-Fernández et al., 2019).

One of the limitations of the ABPP assay is the relatively low throughput due to the complexity of the sample preparation in proteomic applications. Here, we develop methodology to apply activity-based protein profiling in conjunction with enzyme inhibitors in a high-throughput manner, allowing for rapid screening of the concentration dependence and selectivity of multiple inhibitors simultaneously. Although this workflow can be implemented for any ABP containing an affinity tag motif, we used ubiquitin based ABPs to screen cysteine protease deubiquitylating enzymes (DUBs) and a panel of small molecule inhibitors as a methodological proof of concept.

## 2 Materials and Methods

### Cell culture and lysis

MCF-7 cells were cultured in Dulbecco’s Modified Eagle’s Medium (DMEM) with high glucose and supplemented with 10 % (v/v) Fetal Bovine Serum. SH-SY5Y cells were cultured in Eagle’s Minimum Essential Medium and Ham’s F12 Nutrient Mix (1:1), supplemented with 15 % (v/v) Fetal Bovine Serum, 1 % (v/v) non-essential amino acids and 2 mM Glutamax. Cells were maintained at 37 °C, 5 % CO_2_.

For cell collection and lysis, cells were washed with phosphate-buffered saline (PBS), scraped in fresh PBS and collected at 300 xg. Cells were resuspended in lysis buffer (50mM Tris Base, 5 mM MgCl_2_▪6 H_2_O, 0.5 mM EDTA, 250 mM Sucrose, 1 mM Dithiothreitol (DTT), pH 7.5) and vortexed with an equal volume of acid washed glass beads 10 times for 30 seconds, with 2 minute breaks on ice. MCF-7 cells were clarified by 14,000 xg centrifugation at 4 °C for 25 minutes. SH-SY5Y cells were clarified at 600 x G at 4 °C for 10 minutes to retain USP30 bound to mitochondria. Protein concentrations were determined by BCA protein assay.

### DUB small molecule inhibitors used in this study

USP7 inhibitors FT671 and FT827 (Ioannidis et al., 2016; Turnbull et al., 2017) were a kind gift from Stephanos Ioannidis. USP30 inhibitor 39 (Kluge et al., 2018), and USP30 inhibitor 3-b (patent WO2020072964) were kindly provided by Jeff Schkeryantz and Lixin Qiao (Evotec/Bristol-Meyers-Squibb). Inhibitor structures are shown in Figure S3. PR619 was purchased from Calbiochem (Cat. No. 662141), N-Ethylmaleimide from Sigma-Aldrich (Cat. No. E3876), USP7 inhibitor P22077 from Calbiochem (Cat. No. 662142), and USP7 inhibitor HBX41108 from TOCRIS (Cat. No. 4285).

### Tissue collection and lysis

Tissue was harvested from mice culled by exsanguination under terminal anaesthetic (isoflurane >4% in 95%O2 5%CO2); depth of anaesthesia was monitored by respiration rate and withdrawal reflexes. Mice were perfused with PBS and tissue frozen at −80°C. Mouse brain was homogenized in lysis buffer used for the lysis of cultured cells, using a dounce homogenizer. Once the tissue reached a homogenous consistency the glass bead lysis protocol was carried out as outlined for cultured cells. Lysates were clarified at 600 xg at 4 °C for 10 minutes.

### Western blotting

Samples were boiled in Laemmli sample buffer and separated on a Tris-glycine SDS page (4-15 % acrylamide gradient) gel. Samples were then transferred to a PVDF membrane and blocked for 1 h in 4 % milk TBST. Primary antibodies were incubated overnight at 4 °C and secondary antibodies were incubated for 1 hour at room temperature. Imaging was carried out on a LI-COR odyssey detection system.

### Probe synthesis

HA-Ub-PA was synthesized as previously described (Borodovsky et al., 2002; Pinto-Fernández et al., 2019). Ubiquitin was expressed in *E.coli* (Gly76del) with an N-terminal HA tag and a C-terminal intein-chitin binding domain (CBD). *E.coli* were suspended in 50 mM Hepes, 150 mM NaCl, 0.5 mM DTT (buffer used throughout synthesis) and sonicated 10 times, 30 seconds on, 30 seconds off. Purification was carried out using Chitin bead slurry, and HA-Ub-MesNa was formed via overnight agitated incubation with 100 mM MesNa at 37 °C whilst the protein was still attached to the Chitin beads. HA-Ub-PA was formed by incubation of HA-Ub-MesNa with 250 mM PA with agitation at room temperature for 20 minutes. Excess PA was removed via PD-10 column desalting. Complete and active probe formation was confirmed via anti-HA western blot and intact protein LC-MS (data not shown).

### Probe and inhibitor labelling

Lysates were diluted to 3.33 mg/ml using lysis buffer (minus the volume of the inhibitor and probe). Inhibitors were diluted with either DMSO (or ethanol in the case of NEM) to the same volume for their concentration dependence. 3-b, 39, FT671 and NEM were incubated with cell/tissue lysates for 1 hour at 37 °C. Probe was incubated with lysate at a ratio of 1:200 (w/w) for 45 minutes at 37 °C in all conditions, except for FT671 (10 minute incubation) due to long probe incubations displacing bound inhibitor in this case. Reactions were quenched by addition of SDS to 0.4 % (w/v) and NP40 to 0.5 % (v/v) and made to 1 mg/ml protein concentration by addition of NP40 buffer (50 mM Tris, 0.5 % NP40 (v/v), 150 mM NaCl, 20 mM MgCl_2_, pH 7.4) (freezing at this point had no effect on the result of the subsequent IP).

### Immunoprecipitation with Agilent Bravo AssayMAP liquid handling robot

Anti-HA (12CA5) antibody (Roche) was immobilized on Protein A (PA-W) cartridges (Agilent, G5496-60000), using the Immobilization App (Agilent Sample Prep Workbench v3.0.0). All steps use PBS buffer (Sigma-Aldrich). Cartridges were primed with 100 μL (at 300 μL/min) and equilibrated with 50 μL (at 10 μL/min) followed by loading of 100 μg antibody (or otherwise as stated) in a final volume of 50 μL PBS buffer at 3 μL/min. A cup wash with 50 μL and an internal cartridge wash step (100 μL at 10 μL/min) were performed before re-equilibrating the cartridges with 50 μL at 10 μL/min).

The Affinity Purification App was used for pull-downs. Briefly, Protein A cartridges with immobilized anti-HA antibody were primed (100 μL at 300 μL/min) and equilibrated (50 μL at 10 uL/min) with NP-40 buffer, which was also used for all following steps. The sample was loaded at a flow-rate of 1 μL/min. After sample loading the cup was washed (50 μL) and an internal cartridge wash step (100 uL at 10 μL/min) performed to remove unbound lysate. Peptides were eluted using 50 μL at 5 μL/min 6M Urea or 0.15% TFA or 5 % SDS.

### Mass spectrometry sample preparation and analysis

Urea and TFA eluates were diluted/neutralized with 180 μL 100 mM TEAB. Samples were digested in solution with 1 μg Trypsin (Worthington, LS003740 TPCK-treated Trypsin) over night at 37 C. Digestion was stopped by acidification to final concentration of 1% formic acid.

SDS eluates were prepared following an S-Trap 96-well plate (Protifi LLC, C02-96well) protocol. Eluates were acidified with ~12% phosphoric acid (10:1 v/v) and loaded into the S-trap containing 350 μL 90% MeOH in 100 mM TEAB and spun at 1500x g for 1 min. This step was repeated three times. Then samples were resuspended in 100 μL 100 mM TEAB with 1 μg Trypsin (Worthington) and digested over night at 37 C. Samples were eluted from the S-traps in three consecutive steps, each for 1 min at 1500x g, first with 50 μL 50 mM TEAB, then 50 μL 0.1% TFA and finally 50 μL 50% ACN /0.1% TFA. The combined eluates were dried down in a vacuum centrifuge and resuspended in 2% ACN / 0.1% TFA for LC-MS.

### LC-MS data acquisition

Samples were either run on a LC-MS setup comprised of an Evosep One coupled to a Bruker timsTOF Pro or a Dionex Ultimate3000 coupled to a Thermo Q Exactive Classic.

Evotips (Evosep) were prepared and loaded with peptides as described by the manufacturer. Briefly, Evotips were activated by soaking in isopropanol and primed with 20 μL buffer B (ACN, 0.1% FA) by centrifugation for 1 min at 700g. Tips were soaked in isopropanol a second time and equilibrated with 20 μL buffer A (water, 0.1% FA) by centrifugation. 20 μL buffer A were loaded onto the tips and the samples were added. Tips were spun and then washed with 20 μl buffer A followed by overlaying the C18 material in the tips with 100 μL buffer A and a short 20 s spin.

Peptides were separated on an 8 cm analytical C18 column (PepSep, EV-1109, 3μm beads, 100 μm ID) using the pre-set 100 samples per day gradient on the Evosep One. MS data was acquired in PASEF mode (oTOF control 6.2.105 / HyStar 5.1.8.1) in a mass range of 100 −1700 m/z with 4 PASEF frames (3 cycles overlap). The ion mobility window was set from 1/k0 0.85 Vs/cm^2^ to 1.3 Vs/cm^2^, ramp time 100 ms with locked duty cycle.

On the Orbitrap setup comprised of a Dionex Ultimate 3000 nano LC with Thermo Q Exactive Classic peptides were separated on a 50-cm EasySpray column (Thermo Fisher, ES803, 2 μm beads, 75-μm ID) with a 60 minute gradient of 2 to 35% acetonitrile in 0.1% formic acid and 5% DMSO at a flow rate of 250 nL/min.

MS1 spectra were acquired with a resolution of 70,000 and AGC target of 3e6 ions for a maximum injection time of 100 ms. The Top15 most abundant peaks were fragmented after isolation with a mass window of 1.6 Th at a resolution of 17,500 with a maximum injection time of 128 ms. Normalized collision energy was 28% (HCD).

Data for experiment in Figure S1D was generated as follow: Samples of the lysate titration on cartridge have been run on a timsTOF Pro (OtofControl 6.0.115 / HyStar 5.0.37.1) coupled to a Dionex Ultimate 3000 on a 15 cm IonOpticks Aurora series column (1.6 μm beads, 75 μm ID, Ionopticks AUR2-15075C18A) at a flow rate of 400 nL/min. The gradient started for 3 min at 2% B increasing linearly in 17 min to 30% B followed by ramping up to 95% for 1 min and re-equilibration to 2% B. Data has been acquired in PASEF mode as described above.

Data in Figure 5 with compounds FT671, FT827, HBX41108, P22077, 3-b, 39, and PR619 has been acquired on a 100 samples per day gradient on a 8 cm Pepsep column (1.5 μm beads, 150 μm ID, PepSep, PSC-8-150-15-UHP-nC) with PASEF data acquisition as described above.

### Maxquant analysis

Orbitrap raw data was searched in Maxquant 1.6.10.43, timsTOF data was searched in Maxquant 1.6.14. MCF-7 and SHSY5Y cell samples against a reviewed Homo sapiens Uniprot database (retrieved 31-Dec 2018), mouse brain against a reviewed Mus musculus Uniprot database (retrieved 17-Oct 2020).

Maxquant default settings have been used with oxidation of methionine residues and acetylation of the protein N-termini set as variable modifications and carbamidomethylation of cysteine residues as fixed modification. The match between runs feature was used for all analyses.

Raw data and Maxquant search results have been deposited to PRIDE with the identifier PXD023036.

### Data analysis

Graphs were generated and fitted using Graphpad Prism version 8.4.3 (686). All intensities from mass spectrometry experiments are LFQ (label-free quantitative) intensities unless otherwise stated. DUBs were filtered and removed based off presence in a no probe control sample, missing values in the probe control sample, or intensity values that were at the bottom limit of the MS dynamic range.

## 3 Results

### ABBP-HT workflow optimization

Each stage of the high-throughput IP methodology outlined in **Figure 1**, was optimized for maximal DUB identification coverage with minimal material to reduce experimental cost/time. Different starting material type and concentration, probe labeling, immunoaffinity purification, elution, and sample preparation conditions were tested and optimized.

**Figure 1.**
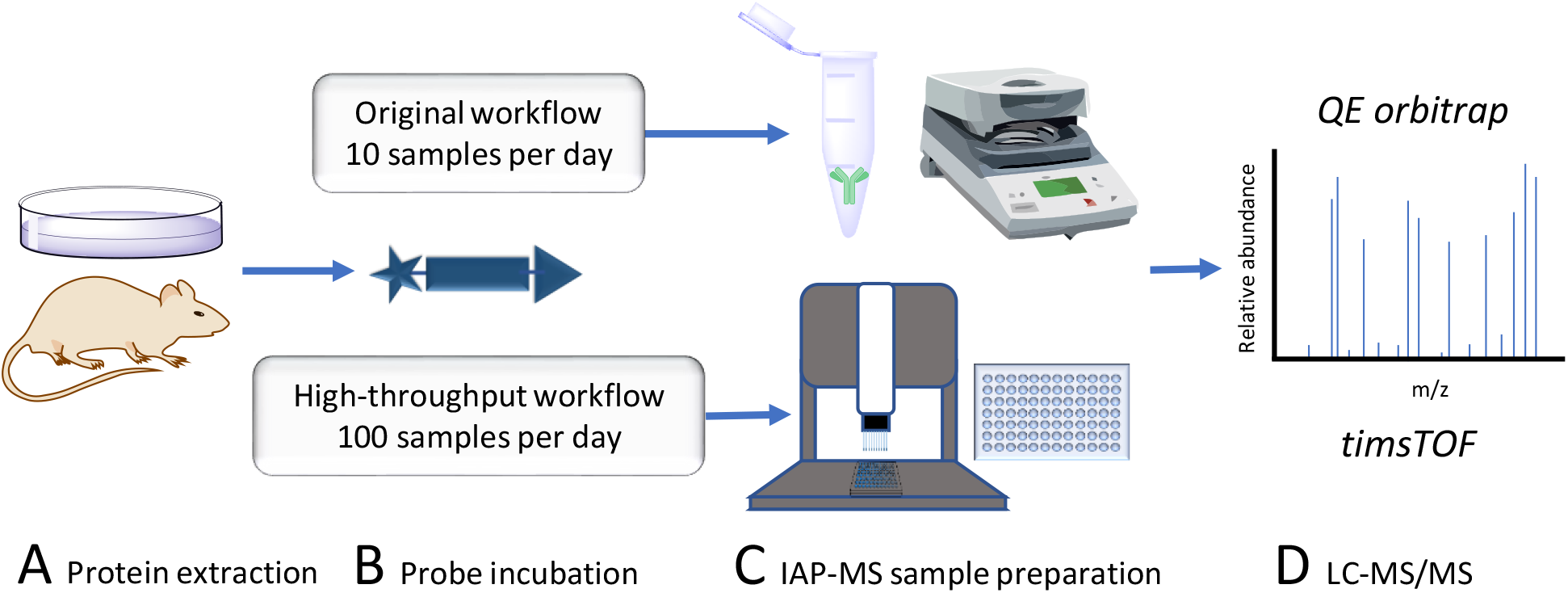
Accelerated DUB inhibitor ABP IP workflow. **A.** Protein extraction and inhibitor treatment of either intact or lysed tissue/cell lines. **B.** HA-Ub-PA probe incubation to label uninhibited cysteine active DUBs. **C.** Anti-HA IP, traditionally with centrifugation or magnetic collection of agarose beads in a low throughput format. In this work the throughput is increased to a 96 well plate format using an Agilent bravo liquid handling platform. **D.** LC-MS/MS proteomic analysis of immunoprecipitated DUBs. Here we compare the depth of the DUBome obtained using a QE orbitrap vs. a high-throughput timsTOF.

**Table 1:**
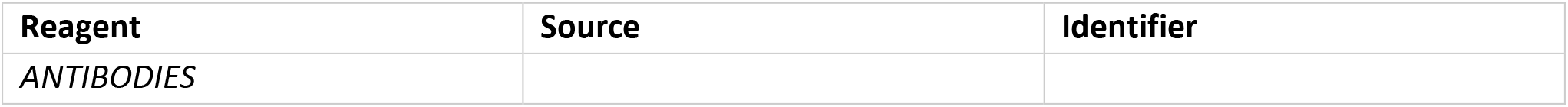
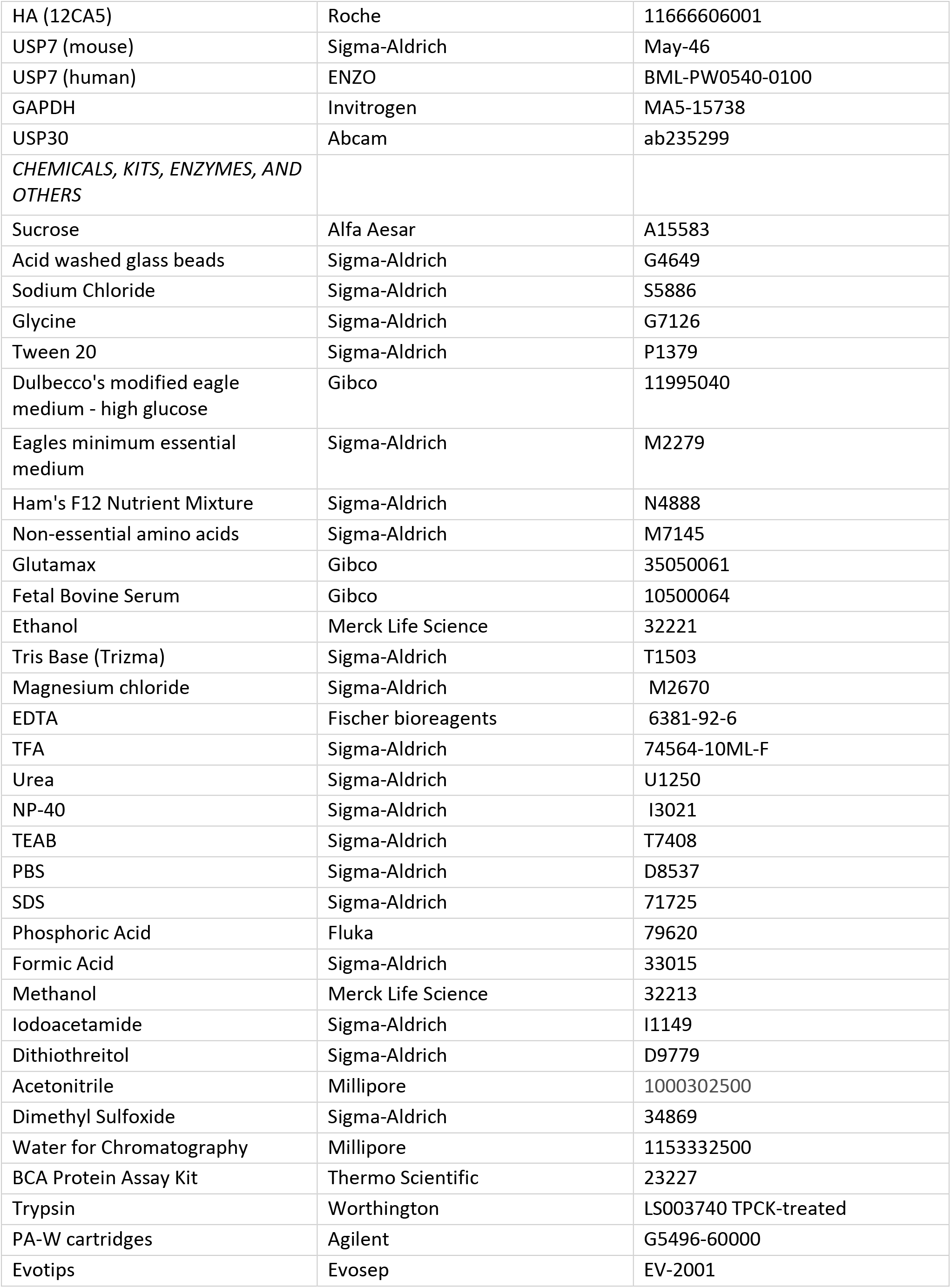
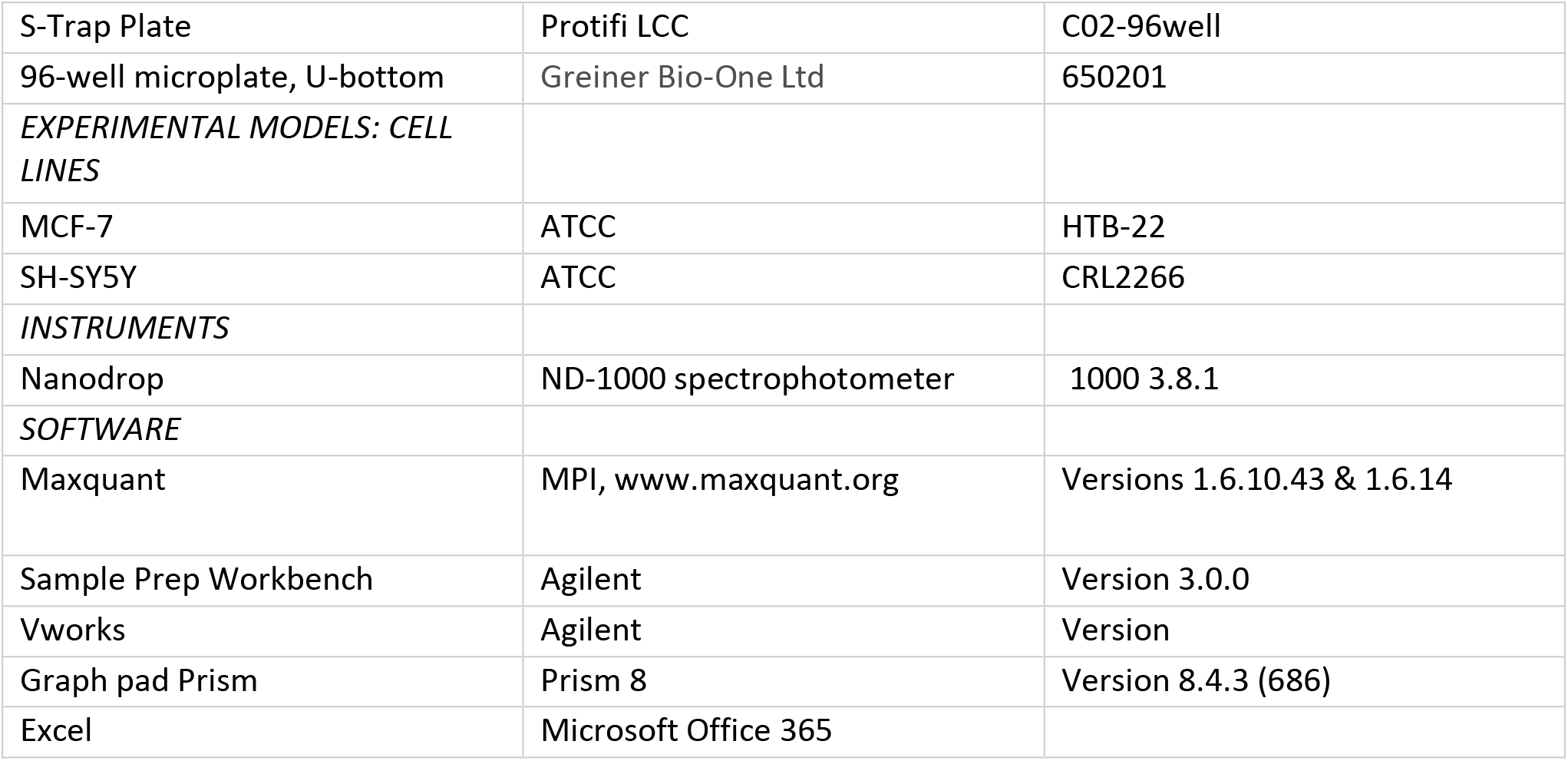
List of reagents.

First, we decided to identify the ideal ratio of probe to protein in the lysate by incubating increasing concentration of the probe with a fixed amount of lysate. Note: this step should be carried out every time after changing the batch of ABP. Our results (shown in **Figure S1A**) suggested that we should use at least 0.25 μg of HA-Ub-PA probe to label efficiently 50 μg protein extracts from two different cell lines, MCF-7 and SH-SY5Y.

In parallel, we also determined the best antibody concentration for immunoprecipitation. Anti-HA antibody was loaded onto Protein A cartridges as a set volume of 50 μl at a flow rate of 3 μg/uL. The column was then washed with 100 μL of PBS buffer. From this, the loading and washing flow through fractions were collected, and unbound antibody was detected by 280 nm Nanodrop measurements. Whilst residual amounts of protein were detected at low concentrations, a significant amount of protein was present when 90 μg was loaded, suggesting the column saturates between 80-90 μg of antibody (**Figure S1B**). From this, we concluded that above 80-90 μg of antibody saturates all column binding sites.

Using cartridges primed with 80 μg of HA antibody, a concentration dependence was also carried out using varying amounts of HA-Ub-PA probe which was diluted to a set volume of 25 μL and loaded at a flowrate of 3 μL/min. The presence of probe in the loading and washing flow-through fractions was detected by anti-HA immunoblotting. Unbound probe was detected both in loading and washing flow-through fractions when ≥ 5 μg of probe was flowed through the column (**Figure S1C**). Therefore, no more than 2-5 μg of probe should be loaded to avoid column saturation and material waste.

In order to identify the optimal amount of labelled lysate we performed an LC-MS/MS analysis after performing IAP-MS with different amounts of labelled lysate, using the parameters described above. The results (**Figure 2A**) showed that 250 μg of labelled lysate gives us the maximum of DUB identifications (IDs). Analysis of the flow through by immunoblotting of USP7 confirmed these results (**Figure 2B**, **Figure S1D,** and **Figure S2**).

**Figure 2.**
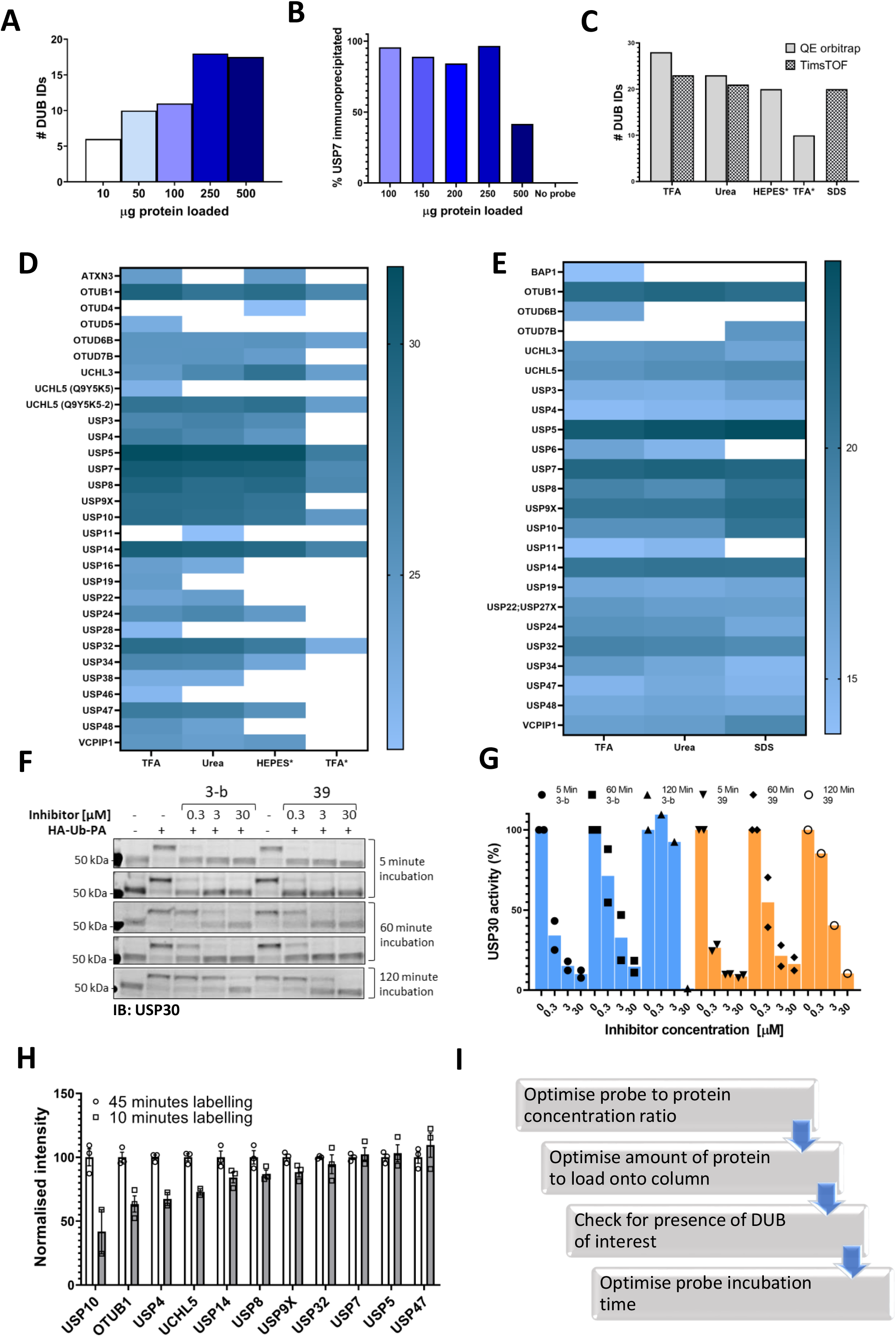
Optimizing the ABPP-HT workflow. **A.** Number of DUBS identified by timsTOF MS with increasing amounts of HA-Ub-PA-labelled MCF-7 lysate protein, after immunoprecipitation and elution with 0.15 % TFA. **B.** Western blot densitometry quantification (full blot in Figure S1D) of USP7 in the immunoprecipitation loading flow-through, with increasing amounts of HA-Ub-PA-labelled MCF-7 lysate protein quantity immunoprecipitated and eluted with 0.15 % TFA. **C.** Number of DUBs identified by LC-MS/MS with different IP elutions: 0.15 % TFA, 6 M urea, HEPES, *= On column digestion. **D.** Log2 intensities of DUBS identified with different elution methods by a QE orbitrap MS. 0.15 % TFA, 6 M urea, HEPES, *= On column digestion. **E.** Log2 intensities of DUBs identified with different elution methods by a timsTOF MS. 0.15 % TFA, 6 M urea, 5 % SDS. **F.** USP30 immunoblots showing mouse brain lysate displacement of a covalent (3-b) and non-covalent (39) USP30 inhibitor with increasing HA-Ub-PA (at 37 °C) incubation times. **G.** The densitometric quantification of Figure 2F from the intensity of the HA-Ub-PA-labelled band, normalised to the intensity of both USP30 bands together. **H.** timsTOF DUB intensities of MCF-7 labelled with HA-Ub-PA for 10 minutes normalised to 45 minutes at 37 °C (SEM, n=3). **I.** Optimisation workflow for high-throughput DUB inhibitor screening using ABPP LC-MS/MS.

The next step was to optimize the elution of the immunoprecipitated material from the column, the digestion method, and the LC-MS/MS instrumentation. Various elution/digestion methods were trialed. Initially 6 M Urea or 0.15 % TFA were used to elute the proteins for digestion and MS analysis. An on-column trypsin digestion of the protein was also carried out in the presence of either HEPES buffer or 0.15 % TFA. Samples were then run on a Q Exactive Orbitrap mass spectrometer and quantified using the search software Maxquant. From this, the most efficient elution was 0.15 % TFA in combination with in-solution trypsin digestion. Different elution methods followed by in-solution digestions were then trialed on an Evosep (liquid chromatography; LC) and timsTOF Pro (mass spectrometry; MS). Comparison of 5 % SDS, 0.15 % TFA and 6M urea using this instrumentation demonstrated that 0.15% TFA is again the most efficient elution for the identification and quantification of the highest number of DUBs (**Figure 2C-E**).

The combined use of short gradients on an Evosep LC and the fast scan speeds using Parallel Accumulation Serial Fragmentation (PASEF) data acquisition on a timsTOF Pro allowed for increased sample throughput. With the preset gradients on the Evosep throughput ranges from 30-300 samples per day (SPD), compared to the 6-12 SPD of common nanoflow LC setups or 9 SPD on our in-house 1 h gradient. In this experimentation 100 SPD is used as standard, with no increase in DUBs identified occurring from a 60 SPD method (data not shown). Additionally, comparison of the Evosep One with a nanoflow LC coupled to the timsTOF Pro resulted in no marked difference in the number of identified DUBs (23 *vs* 22 respectively) on comparable gradient lengths. There is ~ 20% reduction in the number of DUBs identified with the TFA elution using the Evosep/timsTOF compared to an Orbitrap MS (**Figure 2D** and **Figure 2E**) at highly reduced instrument time (single run ~15 min vs 160 min). Both instruments lead to the identification of a similar panel of DUBs, with some DUBs unique to each. The choice between the two instruments should balance the required DUBome coverage *vs* throughput, and whether detection of the desired DUB is feasible with the chosen methodology.

Finally, since one of the applications of this methodology is DUB inhibitor characterization there is an aspect of the ABP assay that should considered before testing a given inhibitor: Probe incubation time. Ub-based activity-based probes, especially with the highly reactive PA (propargylamide) warhead can displace both, covalent and non-covalent, DUB inhibitors over time. As shown in **Figures 2F** and **2G** for two different USP30 inhibitors, the reversible covalent 3-b and the reversible non-covalent 39. Increasing the incubation time displaced the inhibitors, especially 3-b, from the DUB, giving the impression that the inhibitor is less potent. Therefore, for reversible inhibitors we suggest minimizing incubation times with the probe. Of course, this has an impact on the number of DUBs identified when performing the ABPP assay. We compared two labelling times using our ABPP-HT workflow and as expected, the intensities of some DUBs are clearly reduced when the lysate is incubated with probe for a shorter period of time (10 minutes versus 45; **Figure 2 H**). These optimization steps were summarized in **Figure 2I**.

### Guide to DUB picking: abundance changes with methodology and starting material source

These conditions were then applied to characterize the active DUBome in two different cell lines, MCF-7 and SH-SY5Y, and brain tissue material from mice. We also included the data using the two different LC-MS/MS instrumentations: Nanoflow liquid chromatography coupled to an Orbitrap MS (OT on the figures) and microflow (Evosep) liquid chromatography and ion mobility-mass spectrometer, timsTOF (TT on the figures). We summarized the results in a heat map (**Figure 3**), displaying the normalized intensities of the identified DUBs when using different cell lines, tissue, and instrumentation. This together with the heat maps describing the different elution methods (**Figures 2D** and **2E**) should be a good reference when studying a particular DUB and its potential inhibitors. For example, there are some DUBs that are only identified in the cancer cells like USP3, USP4, and OTUD7B. On the other hand, there are DUBs specific for brain tissue and cells like UCHL1.

**Figure 3:**
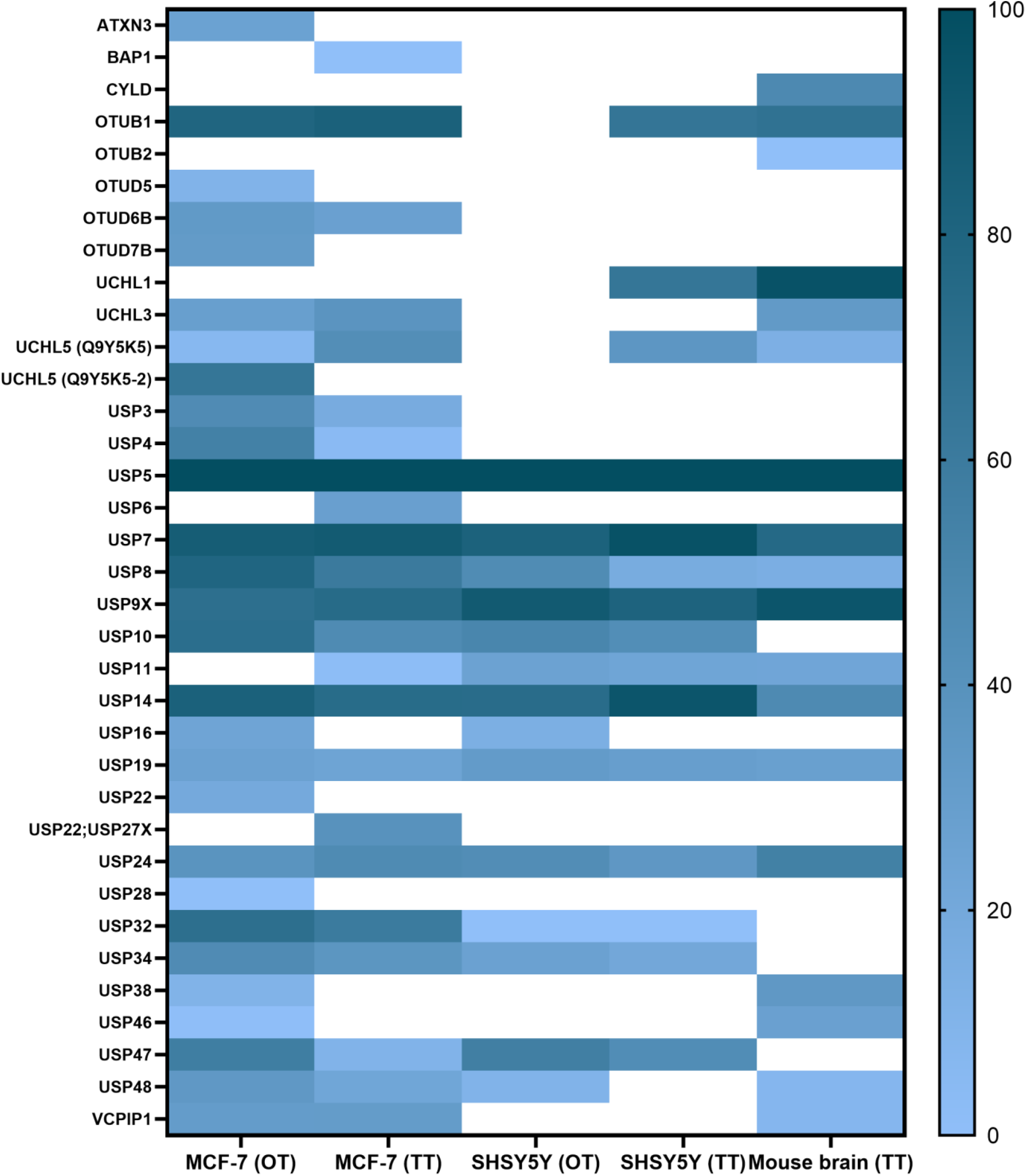
ABPP-HT reveals cell type-specific DUB profiles. DUB intensities as determined by HA-Ub-PA activity-based probe profiling (ABPP) and identified from different cell types and tissue quantified by mass spectrometry using either a QE orbitrap (OT) or timsTOF (TT), normalised within each dataset.

### Proof of concept: Broad and specific DUB inhibitor concentration dependences and cross-reactivities

The ABPP-HT methodology is able to identify a representative panel of DUBs that is comparable to the regular ABPP (~15-25 vs ~30-40) (Pinto-Fernández et al., 2019). With this representative panel we applied the methodology to check for compatibility with DUB inhibitor characterization. In order to do so, we performed ABPP-HT with the highly selective USP7 inhibitor FT671 (Turnbull et al., 2017) and with the broad cysteine modifier NEM (n-ethylmaleimide) (Pinto-Fernández et al., 2019). We performed these experiments treating lysates from MCF-7 cells and from mouse brain tissue extracts with different concentrations of the inhibitors. We used the TT LC-MS/MS instrumentation as it is more suitable for testing a higher number of compounds. First, we performed control immunoblots against USP7, to show the effects of the compounds on USP7, and against the probe (anti-HA), to visualize the selectivity of the compounds. FT671 inhibits USP7 in both, cell line (**Figure 4A**) and brain tissue (**Figure 4B**), however, due to the USP7 antibody recognizing USP7 with and without its previously characterized ubiquitination (Fernández-Montalván et al., 2007), in the mouse tissue this effect was more challenging to visualize. At the same time, the HA blot showed little reactivity of FT671 with other labelled DUBs. On the other hand, NEM was also inhibiting USP7 in both cells (**Figure 4C**) and brain (**Figure 4D**), however, in a non-selective way as shown by the overall decrease in HA signal at high concentrations of the compound. These observations were highly comparable when processing these samples on our ABPP-HT workflow. For instance, immunoblot densitometry of labelled USP7 with increasing amounts of FT671 correlates to a similar degree with the LC-MS/MS data (MCF-7 on **Figure 4E** and brain on **Figure 4G**). This was also the case for the selectivity profile of the two compounds in both cells and brain (**Figures 4F, 4H, 4I,** and **4J**), reflecting well both, the expected high selectivity of FT671 and the broad inhibition by NEM.

**Figure 4.**
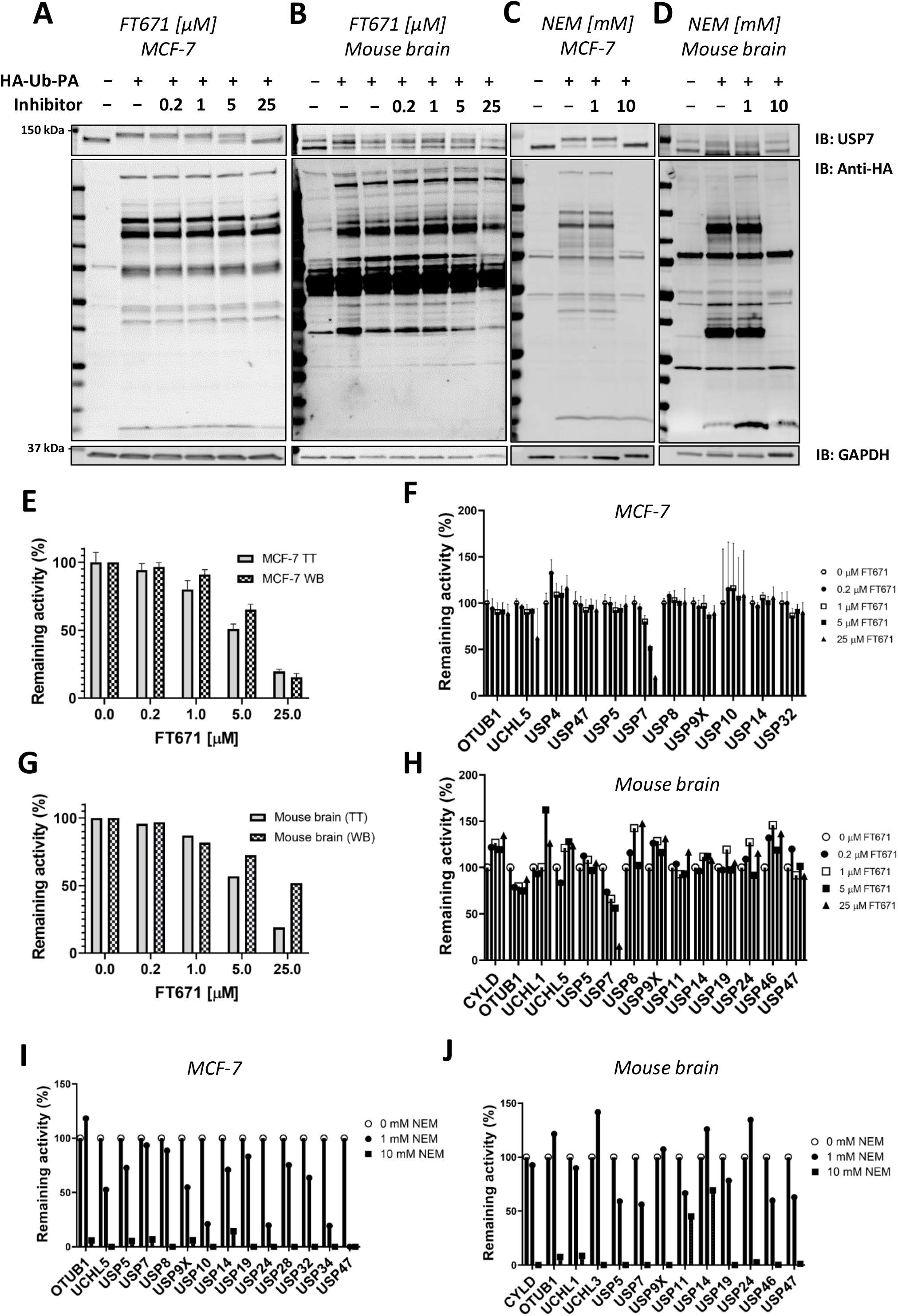
ABPP-HT allows fast generation of DUB inhibitor selectivity profiles in a cellular context. **A-D.** USP7, HA, and GAPDH (loading control) immunoblots of a concentration dependence of FT671 USP7 specific inhibitor and NEM (a general cysteine modifier) in mouse brain tissue and MCF-7 cell lysates. **E.** The densitometric quantification (WB) of three independent experiments as in Figure 4A from the intensity of the HA-Ub-PA USP7 MCF-7 labelled band (SEM, n=3), normalised to the intensity of both USP7 bands together, compared to the MCF-7 timsTOF LFQ normalised intensity (TT) of immunoprecipitated USP7 (MS data SEM n=3 (for 0.2 μM, n=2)). **F.** The activity of a panel of DUBS identified by timsTOF MS from MCF-7 with increasing concentration of FT671 (SEM n=3 (for 0.2 μM, n=2)). **G.** The densitometric quantification of Figure 4B from the intensity of the HA-Ub-PA USP7 mouse brain labelled band, normalised to the intensity of both USP7 bands together (WB), compared to the mouse brain timsTOF LFQ normalised intensity (TT) of immunoprecipitated USP7. **H.** The activity of a panel of DUBS identified by timsTOF MS from mouse brain with increasing concentration of FT671. **I-J.** The activity of a panel of DUBS identified by timsTOF MS from MCF-7 and mouse brain lysates respectively, treated with either 1 mM or 10 mM of NEM.

**Figure 5.**
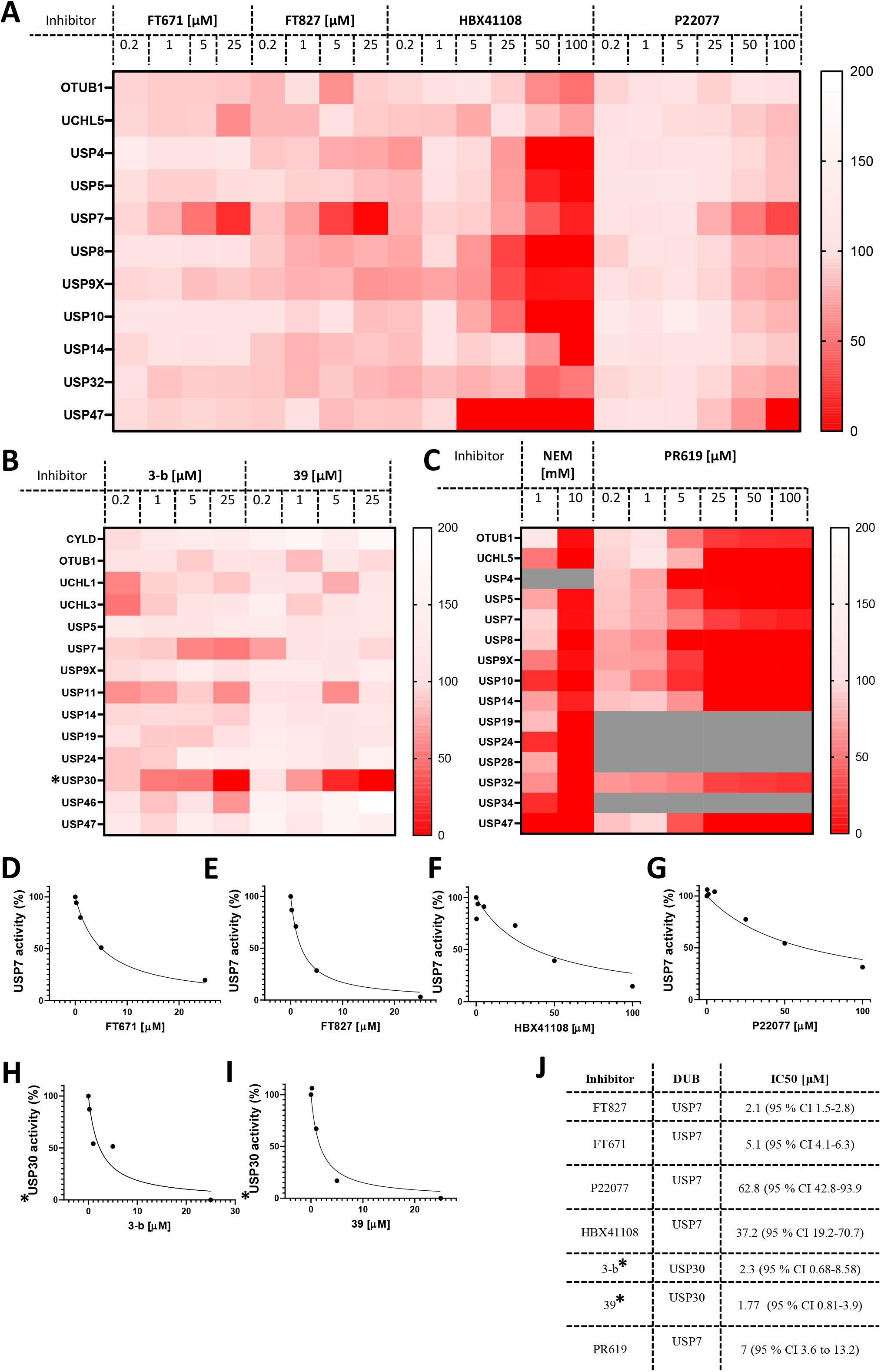
ABPP-HT reveals DUB inhibitor selectivity and specificity compatible with higher throughput. **A.** The activity of a panel of DUBs from MCF-7 identified from timsTOF MS, in response to USP7 specific inhibitors FT671 (n=3 (for 0.2 μM n=2)), FT827, HBX108 and P22077. **B.** The activity of a panel of DUBs from mouse brain lysate identified from timsTOF MS, in response to USP30 specific inhibitors 3-b and 39. **C.** The activity of a panel of DUBs in MCF-7 lysates identified by timsTOF LC-MS/MS, in response to the cysteine modifier NEM, and broad spectrum DUB inhibitor PR619 (PR619 n=2). **D-I.** From left to right concentration dependences of USP7 from inhibitors FT671, FT827, HBX41108, and P22077 in MCF-7 lysates, and USP30 inhibitors for 3-b and 39 in mouse brain. **K.** IC50 values extracted from D-I, fit to equation: Y=100/(1+X/IC50). * = normalised raw intensities, not LFQ intensities.

### Exploiting the possibilities of the ABBP-HT methodology: multiple compound characterization in different cell lines and tissue

Finally, we decided to gain advantage of the ABPP-HT possibilities and applied the methodology to a number of compounds (structures in **Figure S3**) and concentrations simultaneously. Here, critical target engagement information was obtained in a cellular context, in a much faster way than the current methodology. We tested different concentrations of four USP7 inhibitors, FT671, FT827 (Turnbull et al., 2017), HBX41108 (Colland et al., 2009), P22077 (Altun et al., 2011), two USP30 inhibitors (3-b and 39), and the two broad cysteine modifiers NEM (Pinto-Fernández et al., 2019) and PR619 (Altun et al., 2011). The results are summarized on separated heat maps for USP7 inhibitors (**Figure 5A**), USP30 (**Figure 5B**) and non-selective (**Figure 5C**), and bar graphs (**Figures S4A-D**). These results not only match the matching control immunoblots in **Figure S4 (E-H)** but also previously reported information. For instance, P22077 was reported to be a dual USP7/USP47 inhibitor (Altun et al., 2011) and the same result could be seen in our data (Figure 5A). FT671 and FT827 were reported to be highly selective, and potent, USP7 inhibitors (Turnbull et al., 2017) and this still applied when using our ABPP-HT workflow **(Figure 5A)**. HBX41108 selectivity data has not been reported, although our results suggested a not very selective profile. The USP30 inhibitors showed a nice dose-dependent inhibition of the target and good selectivity profiles, especially for the USP30 inhibitor 39 (**Figure 5B**). Finally, the two cysteine modifiers behaved as expected (Altun et al., 2011) (Pinto-Fernández et al., 2019), inhibiting all the identified DUBs at high concentrations (**Figure 5C**). The ability to analyze multiple concentrations also allowed for the plotting and determination of the half-maximal inhibitory concentrations (IC50s) for each inhibitor in the described conditions (**Figure 5D-J**).

## 4 Discussion

There are numerous methods to study the potency and selectivity of an enzyme inhibitor using recombinant purified proteins, and their substrates, in biochemical assays. However, these assays cannot assess the activity of an inhibitor in a more relevant context such as cell lysates, intact cells or tissue. Degradation, limited permeability, or cross-reactivity of an inhibitor in the cellular environment may lead to reduced potency and off-target effects. Consequently, it is important to be able to screen potential inhibitors within this environment. ABPP assays can provide all of these very relevant parameters, however, if we want to apply this technique to a screen of inhibitors with varying concentrations in different cell types, the throughput needs to be increased.

Here, we describe a new ABPP methodology, named ABPP-HT (high-throughput-compatible activity-based protein profiling), that allowed the semi-automated analysis of samples in a microplate format, addressing the low throughput associated to the classic ABPP assay. The incorporation of a liquid handling robot compatible with IAP-MS, and the Evosep/timsTOF LC-MS/MS instrumentation, were key to boost the throughput of the ABPP up to ten times in a cost-effective way. The depth of this method is reduced but comparable to the normal ABPP, with the detection of ~15-25 DUBs when using the ABPP-HT versus ~30-40 DUBs with the original ABPP. The number of DUBs that are reactive with the probe and can be potentially detected by ABPP is higher than 70, but this requires performing a high-pH pre-fractionation of the samples prior LC-MS/MS analysis (Pinto-Fernández et al., 2019). This drastically increases the number of samples to analyze per condition and therefore the required time and cost of the assay. This comparative information has been summarized in **Figure 6**. The methodology of the ABPP-HT approach can be applied as a powerful initial screening tool for multiple inhibitors at different concentrations, in various cell lines, to discard weak or highly cross-reactive inhibitors quickly and robustly. From this, only potent and selective inhibitors could be taken forward for more thorough characterization using the original or fractionated ABPP approaches.

**Figure 6.**
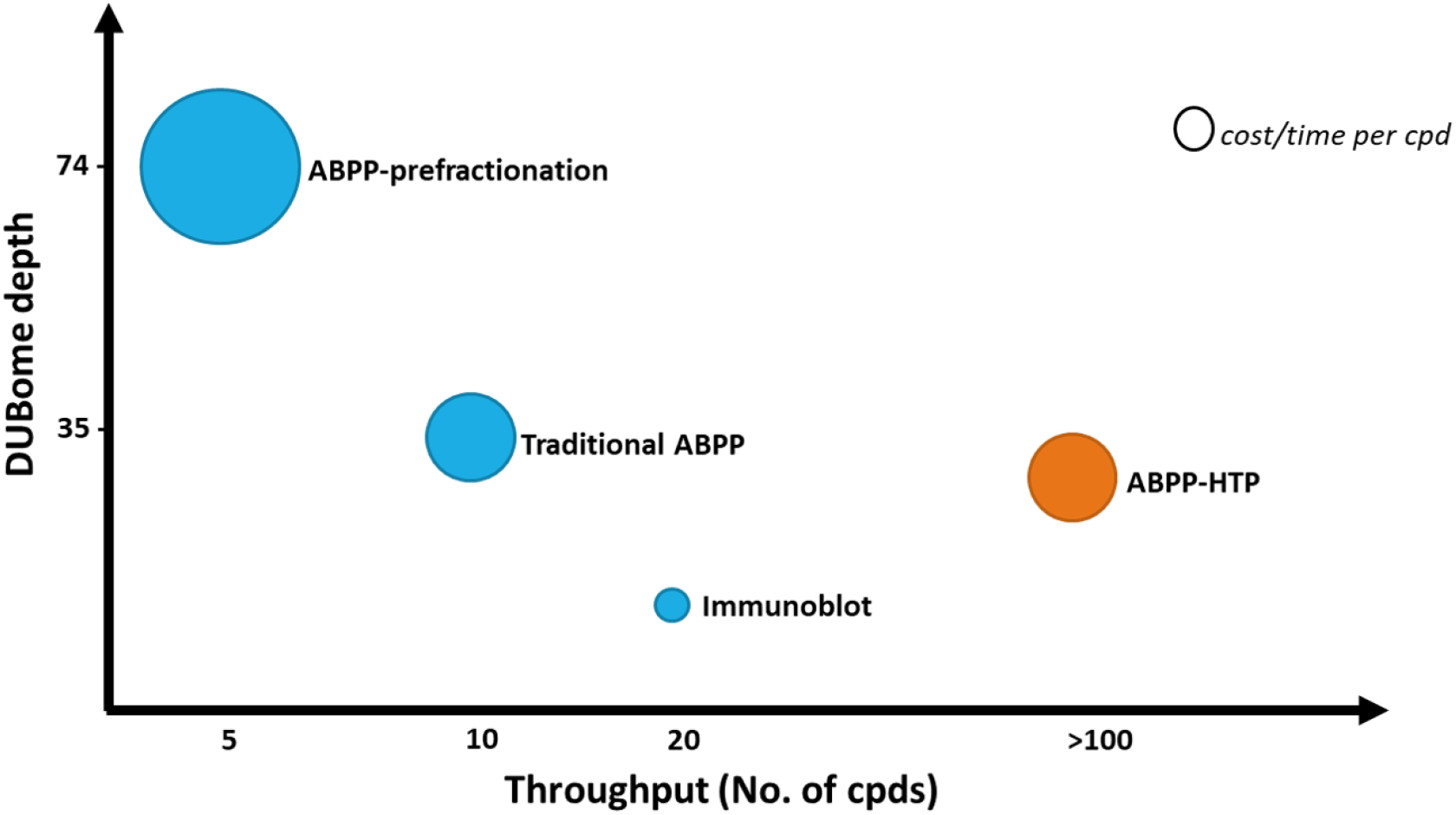
ABPP workflows optimised for DUBome depth and throughput. Comparison of cost and time between ABPP assays optimised for maximal coverage of cellular DUBs (DUBome depth – Y-axis) and higher-throughput approaches including traditional ABPP, immunoblot based ABPP and accelerated throughput ABPP (ABPP-HT) (Throughput – X-axis). Size of the circles are plotted represent relative cost per inhibitor compound tested.

As a proof of concept, we demonstrated the versatility of this methodology using general and specific DUB inhibitors in two different cell lines and mouse brain tissue. ABPP-HT permitted the simultaneous analysis of 6 selective DUB inhibitors and the calculation of their respective half maximal inhibitory concentration (IC50) values. The methodology has the capacity to simultaneously test a much higher number of inhibitors and concentrations. We also believe that the throughput of the ABPP-HT can be increased even further. For example, by implementing chemical labels, such as TMT (tandem mass tag) that would allow the combination of up to 16 samples into one and therefore providing multiplexing capabilities and enhanced throughput. Another area where the sensitivity and therefore DUB coverage of this type of analysis could be further improved is by implementing targeted proteomics methods such as Data-Independent Acquisition (DIA) mass spectrometry. For instance, DIA has been successfully applied for ubiquitomics and discussed by Vere et al. (Vere et al., 2020).

In conclusion, ABPP-HT (high-throughput-compatible activity-based protein profiling) was conceptualized, optimized, and validated. When tested, the approach allowed for reduced time and cost for both sample preparation and MS time, whilst still identifying and quantifying a representative panel of endogenously expressed DUBs, enabling the profiling of a number of DUB inhibitors.

## 6 Author Contributions

H.J., R.F., B.M.K. and A.P.F. directed this study. Most experiments were devised by H.J., R.H., B.M.K. and A.P.F. and carried out by H.J, R.H. and A.P.F. H.J., R.H., B.M.K. and A.P.F. wrote the paper. All authors commented on the text.

## 8 Acknowledgments

We would like to thank the Discovery Proteomics Facility (led by Dr Roman Fischer) at the Target Discovery Institute (Oxford) for expert help with the analysis by mass spectrometry. We would also like to thank Jeffrey M. Schkeryantz (Bristol-Myers Squibb Research and Development), Lixin Qiao (Bristol-Myers Squibb Research and Development), and Katherine England (ARUK Oxford Drug Discovery Institute ODDI) for kindly providing the USP30 inhibitor compounds 39 and 3-b’. Finally, brain tissue material was kindly provided by Gillian Douglas (Division of Cardiovascular Medicine, Radcliffe Department of Medicine, University of Oxford).

## Data Availability Statement

The mass spectrometry proteomics data have been deposited to the ProteomeXchange Consortium via the PRIDE (Perez-Riverol et al., 2019) partner repository with the data set identifier PXD023036.

## Ethical Statement for studies involving animal subjects

The breeding of mice was carried out in accordance with Animal [Scientific Procedures] Act 1986, with procedures reviewed by the University of Oxford clinical medicine animal care and ethical review body (AWERB), and conducted under project licenses PPL P0C27F69A. All procedures conformed to the Directive 2010/63/EU of the European Parliament.

Tissue was harvested from mice culled by exsanguination under terminal anaesthetic (isoflurane >4% in 95%O2 5%CO2); depth of anaesthesia was monitored by respiration rate and withdrawal reflexes. Mice were perfused with PBS and tissue frozen at −80°C.

---

**Figure S1.**
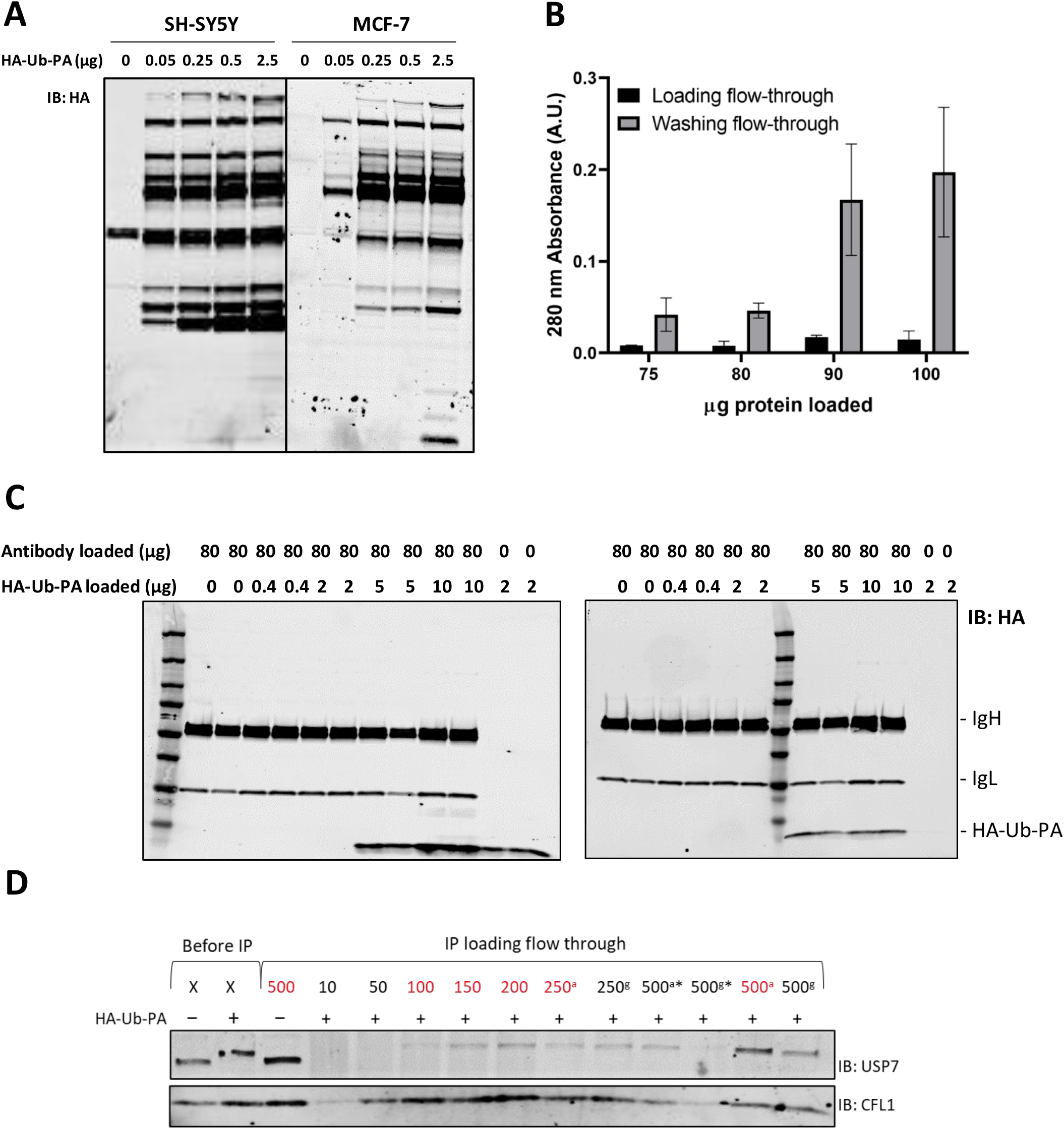
Optimizing DUB target engagement in the ABPP-HT workflow. **A.** HA immunoblot of probe to lysate ratio optimisation, with 50 μg of either SHSY5Y lysate or MCF-7 lysate incubated with increasing amounts of HA-Ub-PA at 37 °C for 45 minutes. **B.** Concentration dependence of unbound antibody detected with antibody loading onto protein A columns. **C.** Concentration dependence of unbound HA-Ub-PA probe detected with HA-Ub-PA loading onto protein G column with 80 μg of antibody loaded. **D.** USP7 detected in IP flow-through where 100 μg of HA antibody has been loaded. Values in red were used for quantitation in figure 2B. ^a^ protein A column, ^g^ protein G column, *material was concentrated before loading to try to minimize loading time, the reduction in material is attributable to protein lost during the concentration step, not to increased HA-Ub-PA anti-HA binding. In each lane 5 ug of protein was loaded except in the case of 10 μg and 50 μg where samples were too dilute to load 5 μg.

**Figure S2.**
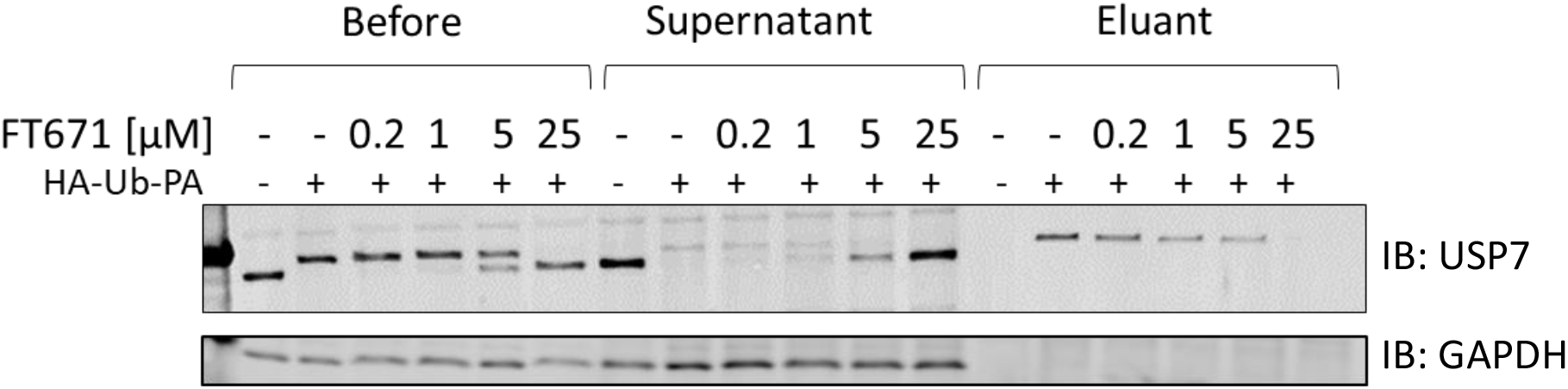
Efficient immunoprecipitation of USP7 with ABPP-HT. USP7 immunoblot of FT671 concentration dependence. Before immunoprecipitation demonstrates probe labelling and inhibition, the supernatant and eluant bands show unlabelled and labelled bands respectively.

**Figure S3.**
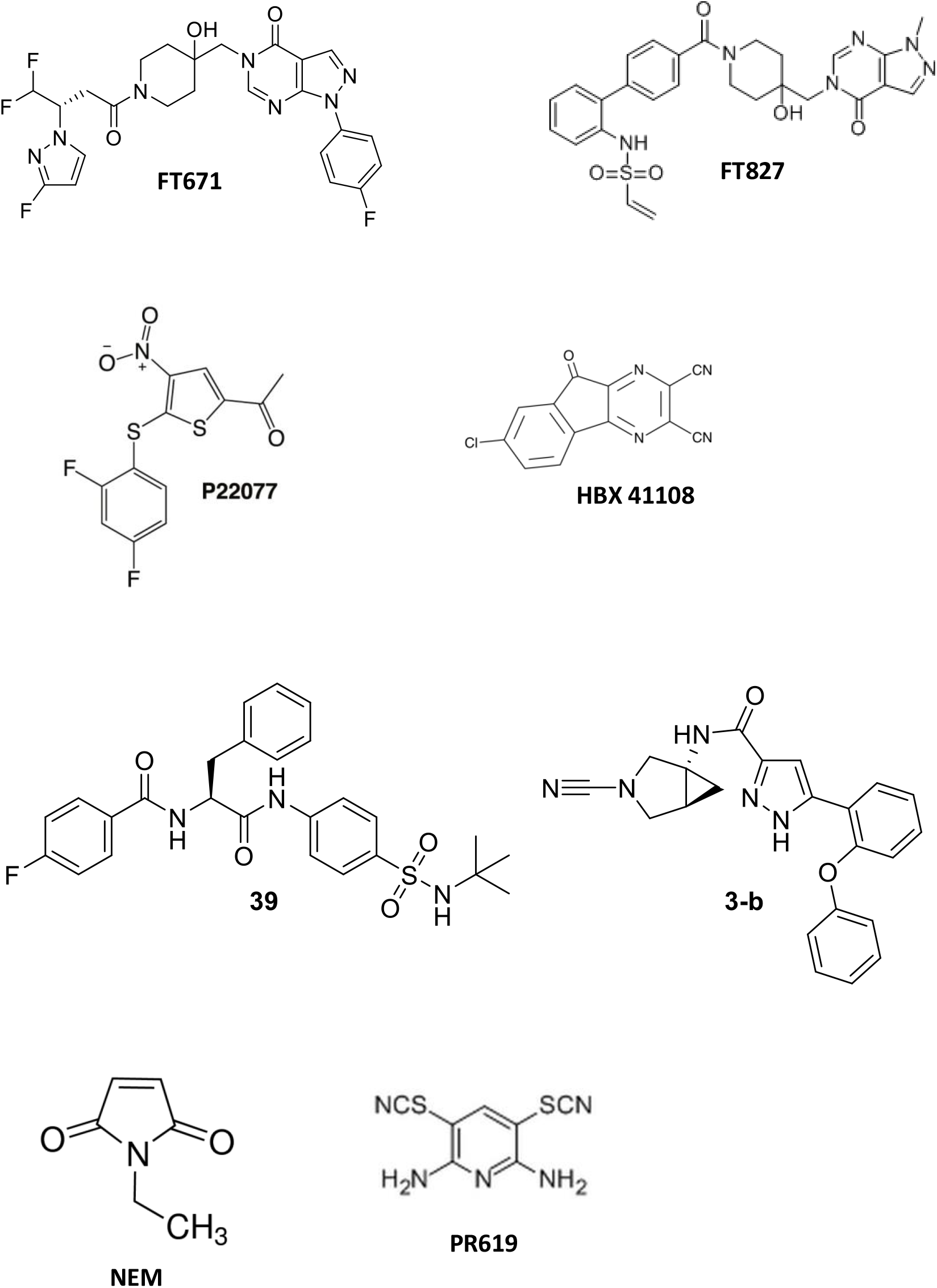
DUB inhibitors used in this study. USP7 inhibitors FT671 FT827, P22077, HBX 41108, USP30 inhibitor 39, USP30 inhibitor 3-b, and broad-spectrum cysteine modifiers NEM and PR619.

**Figure S4.**
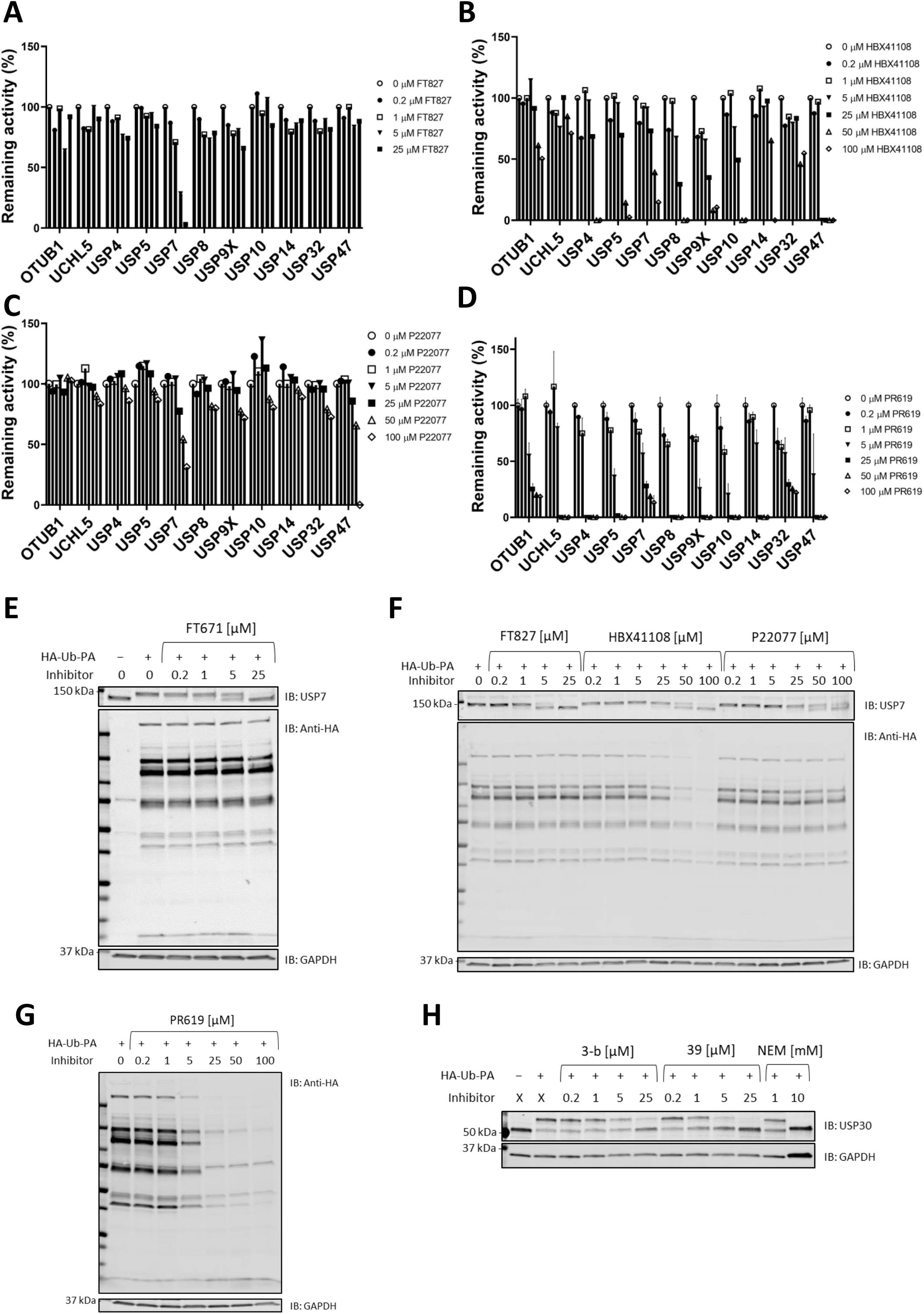
Inhibitor cellular target engagement assessed by ABPP. **A-D.** ABPP inhibition profile of a panel of DUBs by FT827, HBX41108, P22077, and PR609 (one experiment, except for PR609: SEM; n=2). **E.** USP7 and anti-HA immunoblot of specific USP7 inhibitor FT671. **F.** USP7 and anti-HA immunoblot of USP7 inhibitors FT829, HBX108 and P22077. **G.** Anti-HA immunoblot with broad-spectrum DUB inhibitor PR619. **H.** USP30 immunoblot of specific USP30 inhibitors 3-b and 39, and broad DUB inhibitor NEM in mouse brain lysates. GAPDH was used as loading control in these experiments.

